# Lrif1 modulates Trim28-mediated repression of the *Dux* locus in mouse embryonic stem cells

**DOI:** 10.1101/2024.11.18.624083

**Authors:** Darina Šikrová, Román González-Prieto, Alfred C. O. Vertegaal, Judit Balog, Lucia Clemens-Daxinger, Silvère M. van der Maarel

## Abstract

Germline mutations in SMCHD1, DNMT3B and LRIF1 can cause facioscapulohumeral muscular dystrophy type 2 (FSHD2). FSHD is an epigenetic skeletal muscle disorder in which partial failure in heterochromatinization of the D4Z4 macrosatellite repeat causes spurious expression of the repeat-embedded *DUX4* gene in skeletal muscle, ultimately leading to muscle weakness and wasting. All three proteins play a role in chromatin organization and gene silencing; however, their functional relationship has not been fully elucidated. Here, we show that knockdown of *Lrif1*, but not of the other two FSHD2 genes, in mouse embryonic stem cells leads to modest upregulation of the 2-cell cleavage stage transcriptional program driven by the transcription factor Dux, which is the mouse functional homologue of human DUX4. Furthermore, we show that Lrif1 interacts with Trim28, a known Dux repressor and that this interaction is independent of Cbx proteins and Smchd1. We uncover that modest *Dux* upregulation in *Lrif1* knockdown mESCs coincides with decreased Trim28 occupancy at the *Dux* locus. Together, our results provide evidence for a conserved function of Lrif1 in repressing an early zygotic genome activator in mice and humans.

---

Considering the possibility that FSHD is a myo-developmental disease in which a few or all embryonic cells acquire an altered epigenetic landscape of the D4Z4 repeat[1], we tested the potential functional cooperation among FSHD2 gene products in an early developmental cell model. We cultured E14 mouse embryonic stem cells (mESC) in serum/LIF condition where all three FSHD2 (MIM: 158901) genes are co-expressed, and we performed siRNA-mediated knockdown of each of them followed by total RNA-seq (see Supplemental methods for experimental details). In contrast to *Smchd1* (MIM: 614982), for which only a single mRNA isoform is expressed in this culture system, *Dnmt3b* (MIM: 602900) and *Lrif1* (MIM: 615354) give rise to at least three different protein-coding isoforms in E14 mESCs (Figure 1A). Therefore, we used a mix of four siRNAs for each gene to ensure targeting all known isoforms. Cells were harvested after 48 hours of knockdown, which resulted in efficient protein (Figure 1A) and mRNA (Figure 1B) depletion, while mRNA and protein levels of untargeted FSHD2 genes remained unaffected (Figure 1A, 1B). The mRNA levels of pluripotency markers *Oct4* (MIM: 164177) and *Nanog* (MIM: 607937) were largely unchanged upon their respective knockdowns. However, we detected a mild but significant downregulation of *Sox2* (MIM: 184429) mRNA levels upon Lrif1 depletion (Figure 1C). Therefore, short-term depletion of these chromatin factors does not interfere with the expression of tested pluripotency markers in serum/LIF condition. To better understand the roles of these factors in gene regulation, we first performed total RNA-seq. This did not reveal robust expression changes in any of the knockdown conditions (considering expression changes in either direction by at least a factor of 2), with Lrif1 knockdown showing the most differentially expressed genes (749 DEGs with padj <0.05), the majority being modest fold changes (Figure 1D-1F). Despite statistically significant overlaps between differentially upregulated and downregulated genes in paired comparisons of the knockdown conditions, only a limited number of upregulated and downregulated genes (23 and 9, respectively) was shared among all three knockdown conditions (Figure 1G and 1H). This limited overlap prohibits pathway analysis and suggests rather divergent effects of these proteins on the transcriptome in mESCs.

**Figure 1.**
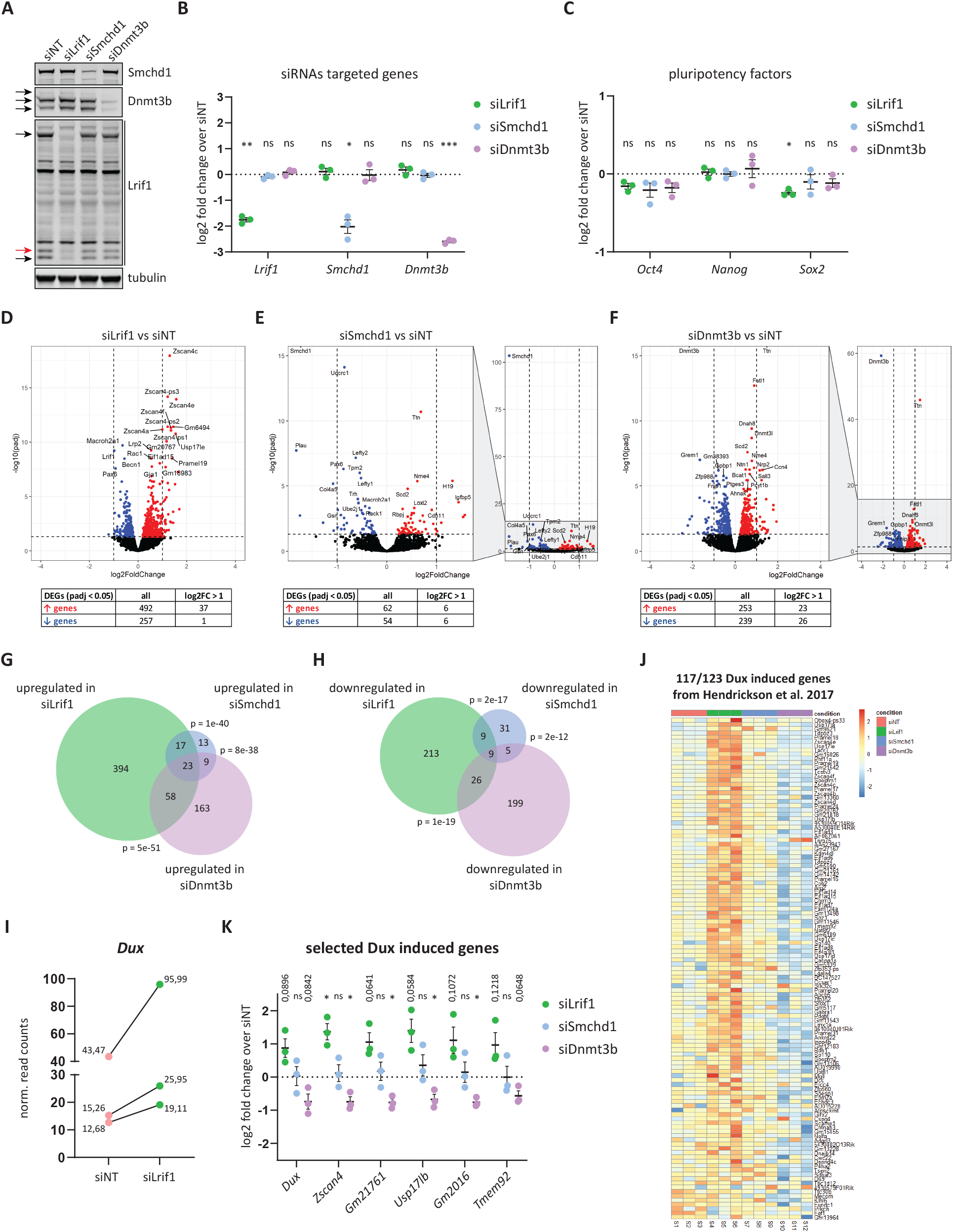
Lrif1 knockdown causes upregulation of Dux sensitive genes. **A)** Western blot confirmation of successful siRNA-mediated knockdown of three FSHD2 genes (*Lrif1, Smchd1* and *Dnmt3b*) in E14 mESCs. Arrows mark different protein isoforms of Dnmt3b and Lrif1. Tubulin served as a loading control. A representative western blot is shown. **B)** RT-qPCR confirmation of all three FSHD2 genes after siRNA-mediated knockdown in E14 mESCs. Expression levels detected in the knockdown conditions were normalized to the siNT condition, and log2 transformed. Each dot represents an independent biological replicate. Whiskers represent mean ± SEM. Statistical significance was calculated by one-sample t-test (ns: not significant, *: < 0.05, **: < 0.01, ***: < 0.001). **C)** RT-qPCR of three pluripotency genes (*Oct4, Nanog* and *Sox2*) after siRNA-mediated knockdown of all three FSHD2 genes. Expression levels in the knockdown conditions were normalized to the siNT condition, and log2 transformed. Each dot represents an independent biological replicate. Whiskers represent mean ± SEM. Statistical significance was calculated by one-sample t-test (ns: not significant, *: < 0.05). **D)** Volcano plot showing gene expression changes following Lrif1 knockdown. Upregulated genes are highlighted in red, and downregulated genes are highlighted in blue. Dashed lines indicate a fold change of two (log_2_fold of 1) on the x-axis and a significance of 0.05 (−log10 p.adj of 1.3) on the y-axis. The top 20 differentially expressed genes (DEGs) are labelled. A table summary of DEGs is provided below the plot. **E)** Volcano plot showing gene expression changes following Smchd1 knockdown. Upregulated genes are highlighted in red, and downregulated genes are highlighted in blue. Dashed lines indicate a fold change of two (log_2_fold of 1) on the x-axis and significance of 0.05 (−log10 p.adj of 1.3) on the y-axis. The top 20 differentially expressed genes (DEGs) are labelled. A table summary of DEGs is provided below the plot. **F)** Volcano plot showing gene expression changes following Dnmt3b knockdown. Upregulated genes are highlighted in red, and downregulated genes are highlighted in blue. Dashed lines indicate a fold change of two (log_2_fold of 1) on the x-axis and significance of 0.05 (−log10 p.adj of 1.3) on the y-axis. The top 20 differentially expressed genes (DEGs) are labelled. A table summary of DEGs is provided below the plot. **G)** Overlap of common significantly upregulated genes (padj < 0.05) in all three knockdown conditions. **H)** Overlap of common significantly downregulated genes (padj < 0.05) in all three knockdown conditions. **I)** Changes in normalized read counts of Dux transcripts after Lrif1 knockdown compared to the non-targeting siRNA condition. **J)** Heatmap of Z-score values depicting expression changes of previously reported 117 Dux-induced genes (out of 123 reported by Hendrickson et al.[8] in the different knockdown conditions for which there was a non-zero read count. **K)** RT-qPCR of Dux and five of its target genes after siRNA-mediated knockdown in E14 mESCs. Expression levels detected in the knockdown conditions were normalized to the siNT condition, and log2 transformed. Each dot represents an independent biological replicate. Whiskers represent mean ± SEM. Statistical significance was calculated by one-sample t-test (ns: not significant, *: < 0.05).

Interestingly, the top differentially upregulated genes after *Lrif1* knockdown (such as the *Zscan* gene cluster) belong to a class of genes which are specifically expressed at the two-cell (2C) cleavage stage of the mouse embryo[2-5] (Figure 1D). The 2C-like cells spontaneously arise in mESC culture, accounting for less than 1% of the population[4], and they mimic some of the distinctive features of the 2C-stage embryos (reviewed here[6]). Furthermore, expression analysis of repetitive elements in *Lrif1* knockdown mESCs showed a significant increase in transcripts originating from repeats, which are known to be de-repressed in the 2-cell embryo and the 2C-like mESCs population such as major satellites[7] and MERVL elements[4] (Figure S1A).

The expression of many of these genes and repeats is driven by the transcription factor Dux, a functional homologue of the primate *DUX4* (MIM: 606009) gene product. *Dux* itself was not significantly differentially upregulated in *Lrif1* knockdown mESCs, however, plotting the normalized read counts of *Dux* in each *Lrif1* knockdown experiment showed a modest increase in read numbers in each knockdown experiment (Figure 1I), whereas *Dux* normalized read counts did not increase in the other two knockdown conditions (Figure S1B and S1C). Therefore, we examined the expression levels of selected genes previously described by Hendrickson et al.[8], which are sensitive to Dux overexpression in mESCs (hereafter referred to as Dux signature genes). This revealed that, in general, the mRNA levels of Dux signature genes are increased upon *Lrif1* knockdown (Figure 1J). In contrast, Dux signature genes remained unchanged in the Smchd1 knockdown situation (Figure 1J). In addition, Dnmt3b knockdown decreased the expression of *Dux* and Dux signature genes (Figure S1C and Figure 1J). We validated with RT-qPCR the mRNA expression of five selected Dux signature genes (*Zscan4, Gm21761, Usp17lb, Gm2016, Tmem92*) and *Dux* itself, and confirmed their upregulation in *Lrif1* knockdown mESCs, albeit not always reaching statistical significance (Figure 1K). We further confirmed the absence of expression effects for these genes in *Smchd1* knockdown mESCs and their mild downregulation upon Dnmt3b knockdown (Figure 1K). Consistent with the RNA-seq, the expression changes were subtle. Therefore, these results indicate that Lrif1 confers a mild repression of the Dux-driven 2C-like transcriptional program in mESCs under these experimental conditions.

Several chromatin-regulating factors have been reported to directly influence *Dux* expression in mESCs[9-13]. Therefore, we decided to investigate the protein interactome of Lrif1 in mESCs to identify potential interactors that could explain Lrif1’s contribution to the regulation of 2C-like cells. To this end, we generated constructs encoding two GFP-tagged Lrif1 isoforms corresponding to the amino acid sequences of the human long (GFP-Lrif1l) and short LRIF1 isoforms (GFP-Lrifs). After transient expression of the Lrif1 fusion proteins in E14 mESCs, we performed a GFP-specific pull-down followed by mass spectrometry (MS). MS analysis identified 54 proteins enriched in GFP-Lrif1s pull-down, of which 37 were nuclear (Figure 2A) and 94 proteins enriched in GFP-Lrif1l pull-down, of which 44 were nuclear (Figure 2B). Since Lrif1 mainly localizes to the nucleus, we focused our further analysis on nuclear proteins[14].

**Figure 2.**
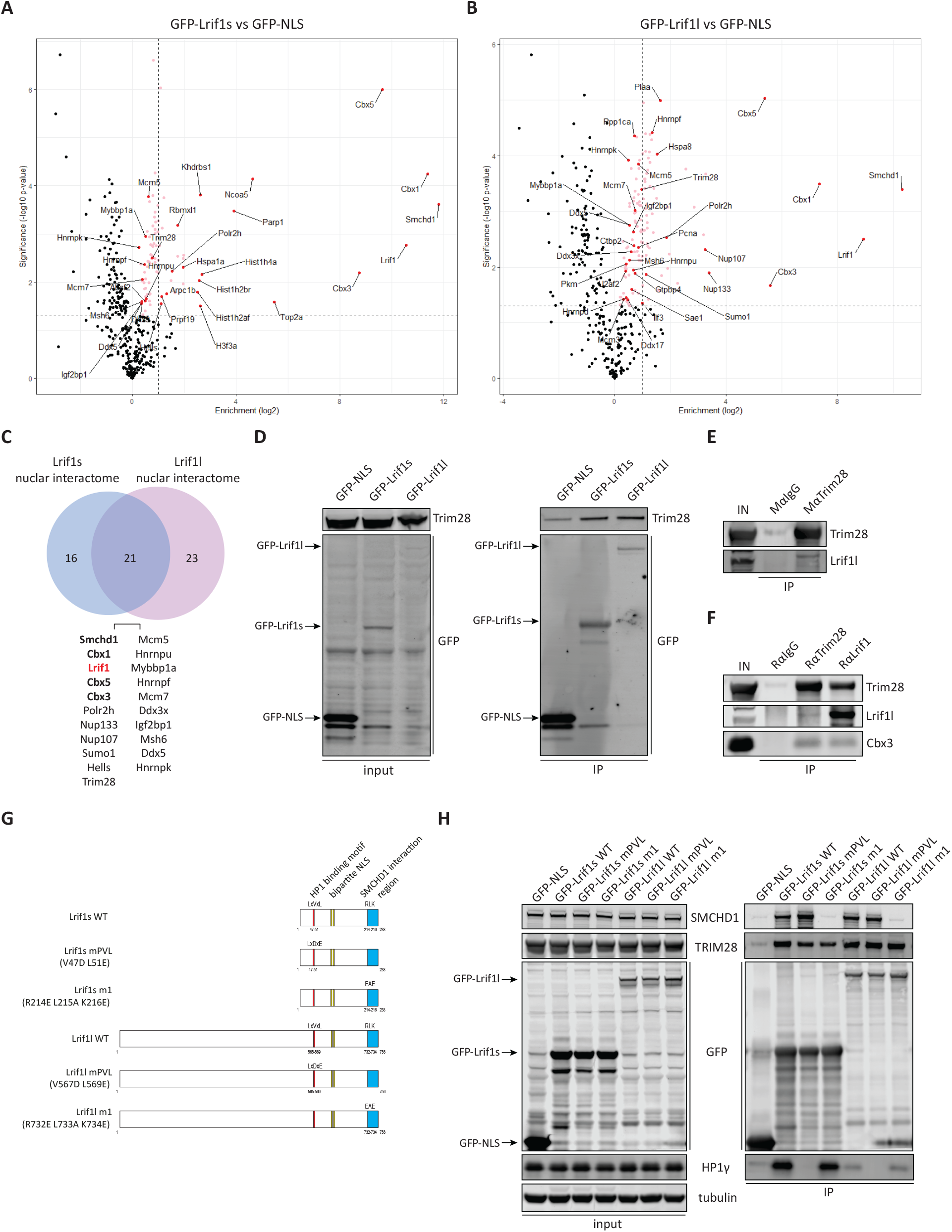
Lrif1 isoforms interact with Trim28. **A)** Volcano plot showing the differential interactome of the GFP-tagged Lrif1 short isoform over GFP-NLS only as identified by GFP pull-down followed by label-free MS. The enrichment (log2) is plotted on the x-axis, and the significance (−log10 p-value) is plotted on the y-axis. The dashed line indicates a significance of 0.05 (−log10 P value of 1.3) on the y-axis. Red dots mark significantly enriched nuclear proteins, and pink dots mark significantly enriched non-nuclear proteins. Ribosomal proteins are not shown. All significantly enriched nuclear proteins are labelled. **B)** Volcano plot showing the differential interactome of the GFP-tagged Lrif1 long isoform over GFP-NLS. The same description applies to this plot as for panel A. **C)** Venn diagram of overlapping significantly enriched nuclear proteins between the GFP-tagged Lrif1 short and long isoform interactomes. **D)** Western blot confirmation of the Lrif1-Trim28 interaction by GFP pull-down of GFP-NLS, GFP-Lrif1s or GFP-Lrif1l. **E)** Endogenous MαTrim28 co-IP on benzonase-treated mESC whole cell extracts. MαIgG was used as a negative control. Only the long isoform of Lrif1 is probed for as the short Lrif1 isoform protein migrates at the height of the IgG light chain. **F)** Reciprocal endogenous RαTrim28 and RαLrif1 co-IP on benzonase-treated mESC whole cell extracts. MαIgG was used as a negative control. Only the long isoform of Lrif1 is probed for as the short Lrif1 isoform protein migrates at the height of the IgG light chain. **G)** Schematic representation of wild-type GFP tagged Lrif1 constructs and their mutant forms used for GFP Co-IPs. **H)** GFP pull-downs of GFP-NLS, wild-type and mutant GFP-tagged Lrif1 isoforms in benzonase-treated HEK293T whole cell extracts.

We found that nuclear proteins enriched in the GFP-Lrif1s pull-down largely overlapped with proteins identified in the GFP-Lrif1l pull-down (Figure 2C). Smchd1 and three Cbx paralogues were among the top four interactors, consistent with previous findings[15-17]. We identified Trim28 (tripartite motif-containing protein 28, also known as Kap1; MIM: 601742) as a common interacting partner of both Lrif1 isoforms. Evidence shows that Trim28 is directly involved in repressing the *Dux* locus in mESCs [9, 12]. In addition, several studies showed that depletion of Trim28 in mESCs cells leads to an increase in the 2C-like population in a Dux-dependent manner[4, 9, 12]. Therefore, to address putative cooperation between Lrif1 and Trim28, we first confirmed the Trim28 interaction with both Lrif1 isoforms by transfecting the GFP-tagged Lrif1 isoforms in mESCs followed by GFP-specific pull-down and western blot analysis (Figure 2D). We further validated this interaction by performing reciprocal endogenous co-Ips from mESC whole-cell extract treated with benzonase to rule out possible DNA-mediated interactions using two different Trim28 antibodies and one Lrif1 antibody (Figure 2E and 2F). Detection of endogenous Lrif1s was hindered by its co-migration with the antibody light chain used for co-IP, which was of the same species origin as the primary antibody used for Lrif1 detection.

We could also pull-down Cbx3 (MIM: 604477) in the Trim28 co-IP (Figure 2F) in agreement with previously published observations[18]. Like Lrif1, Trim28 contains a conserved Cbx binding motif (PxVxL; x represents any amino acid), essential for transcriptional silencing imposed by Trim28[19]. We speculated that the interaction between Lrif1 and Trim28 is mediated via Cbx proteins, the homologues of human HP1 proteins[20]. To test this hypothesis, we introduced previously characterized mutations (mPVL) in the HP1 binding motif of the human LRIF1 [17] to our GFP-tagged long and short mouse Lrif1 constructs (Figure 2G). The mPVL mutant carries two amino acid substitutions (V47D/L51E in the short isoform; V567D/L569E in the long isoform) in the conserved HP1 binding motif, which abolish the interaction of LRIF1 with the chromoshadow domain of HP1 proteins. We included an additional LRIF1 mutant termed m1[17] in this experiment. This mutant carries three amino acid substitutions in the C-terminal coiled-coil domain (R214E/L215A/K216E in the short isoform; R732E/L733A/K734E in the long isoform) of LRIF1, which compromises the interaction with SMCHD1[17]. We transfected mouse mPVL and m1 mutant constructs into HEK293T cells and performed GFP-specific pull-down and western blot analysis. As anticipated due to functional motif conservation[17], both mouse Lrif1 isoforms interacted with the human SMCHD1 and CBX3/HP1γ proteins as well as with human TRIM28 (Figure 2H). Next, we assessed the Trim28 interaction with the mutated forms of Lrif1. As expected, the mPVL mutations abolished the interaction of Lrif1 with human HP1γ, which corresponds with mouse Cbx3, and the m1 mutant of Lrif1 failed to interact with human SMCHD1. Surprisingly, neither of the mutants affected Lrif1’s interaction with TRIM28 (Figure 2H). This suggests that the interaction of Lrif1 with TRIM28 is not mediated via HP1 proteins nor via SMCHD1 or its coiled-coil domain and that another region shared by both Lrif1 isoforms is responsible for this interaction.

Since evidence was presented that Trim28 represses *Dux* by directly binding to its genomic locus[9, 12] and we uncovered an interaction between Lrif1 and Trim28, we investigated a potential interplay of Lrif1 and Trim28 at the *Dux* locus. First, we employed siRNA-mediated short-term depletion of either Lrif1 or Trim28 and confirmed their knockdown efficiency by western blot (Figure 3A) and RT-qPCR (Figure 3B). Lrif1 knockdown did not affect protein levels of Trim28 or the other two Lrif1 interacting partners (Smchd1 and Cbx3), also known to regulate *Dux*[10, 21]. This result rules out the possibility that the observed increased *Dux* expression in Lrif1 knockdown is due to lower levels of any of these *Dux* repressors. A two day-long knockdown of Trim28 was already sufficient to cause mild downregulation in the expression of three examined pluripotency factors (Figure 3C), which is in agreement with its essential role in pluripotency maintenance and self-renewal of mESCs cultured in serum/LIF condition[22]. Despite that, we could detect a modest upregulation of *Dux* and five of its signature genes by RT-qPCR in both knockdown situations (Figure 3D).

**Figure 3.**
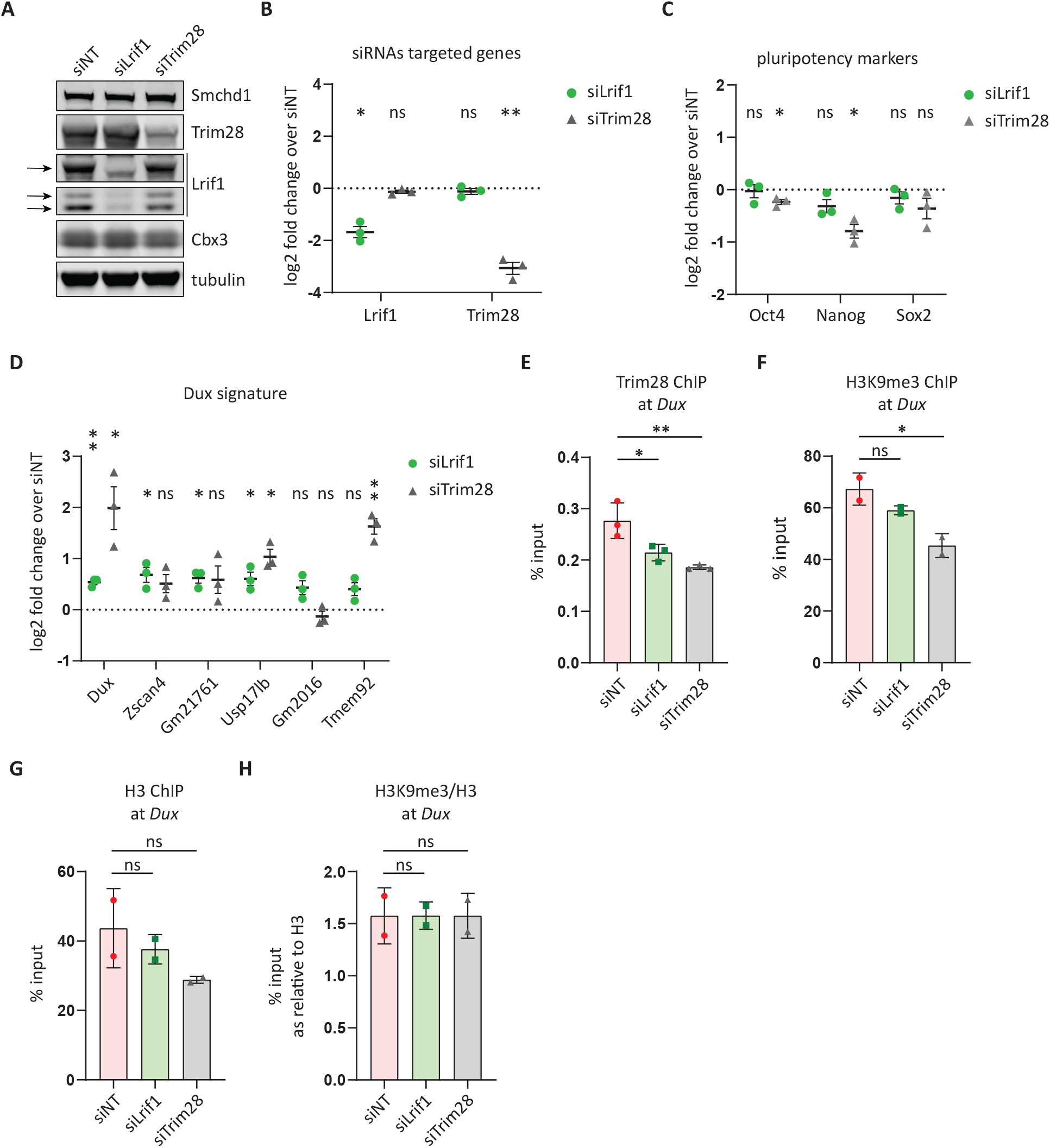
Trim28 requires Lrif1 for its binding to *Dux* repeat. **A)** Western blot confirmation of successful siRNA-mediated knockdown of Lrif1 or Trim28 in E14 mESCs. Arrows mark different protein isoforms of Lrif1. Tubulin served as a loading control. The protein levels of the other two Lrif1 interactors (Smchd1 and Cbx3) are unchanged. A representative western blot is shown. **B)** RT-qPCR confirmation of *Lrif1* and *Trim28* downregulation after siRNA-mediated knockdown in E14 mESCs. Expression levels detected in the knockdown conditions were normalized to the siNT condition, and log2 transformed. Each dot represents an independent biological replicate. Whiskers represent mean ± SEM. Statistical significance was calculated by one-sample t-test (ns: not significant, *: < 0.05, **: < 0.01). **C)** RT-qPCR of three pluripotency genes (*Oct4, Nanog* and *Sox2*) after siRNA-mediated knockdown of Lrif1 or Trim28. Expression levels in the knockdown conditions were normalized to siNT condition, and log2 transformed. Each dot represents an independent biological replicate. Whiskers represent mean ± SEM. Statistical significance was calculated by one-sample t-test (ns: not significant, *: < 0.05). **D)** RT-qPCR of *Dux* and five of its target genes after siRNA-mediated knockdown of Lrif1 or Trim28. Expression levels in the knockdown conditions were normalized to the siNT condition, and log2 transformed. Each dot represents an independent biological replicate. Whiskers represent mean ± SEM. Statistical significance was calculated by one-sample t-test (ns: not significant, *: < 0.05, **: < 0.01). **E)** Trim28 ChIP-qPCR of the 5’ *Dux* region in E14 mESCs after treatment with respective siRNAs. Bars and whiskers represent mean ± SEM of three independent experiments. Statistical significance was calculated by one-way ANOVA with ‘Dunnett’s post hoc test (*: < 0.05, **: < 0.01). **F)** H3K9me3 ChIP-qPCR of the 5’ *Dux* region in E14 mESCs after treatment with respective siRNAs. Bars and whiskers represent mean ± SEM of two independent experiments. Statistical significance was calculated by one-way ANOVA with ‘Dunnett’s post hoc test (ns: not significant, *: < 0.05). **G)** H3 ChIP-qPCR of the 5’ *Dux* region in E14 mESCs after treatment with respective siRNAs. Bars and whiskers represent mean ± SEM of two independent experiments. Statistical significance was calculated by one-way ANOVA with Dunnett’s post hoc test (ns: not significant). **H)** Ratio of H3K9me3 levels to H3 levels at *Dux* calculated from enrichment values presented in F) for H3K9me3 and G) for H3. Statistical significance was calculated by one-way ANOVA with Dunnett’s post hoc test (ns: not significant).

Next, we wanted to assess if Lrif1 directly regulates the *Dux* locus by chromatin immunoprecipitation (ChIP). Since no validated ChIP-grade antibody for mouse Lrif1 is available, we focused on a potential Lrif1-dependent Trim28 binding to the *Dux* locus. As expected, Trim28 knockdown leads to decreased Trim28 enrichment at the *Dux* locus (Figure 3E) as well as at IAPEz elements (Figure S3A), which are also innate genomic targets of Trim28-imposed repression in mESCs[22, 23]. Interestingly, Lrif1 knockdown reduced Trim28 enrichment at *Dux*, albeit less than observed in the Trim28 knockdown condition (Figure 3E). In contrast, the Trim28 enrichment at IAPez elements remained unaffected in Lrif1 knockdown mESCs (Figure S3A) in agreement with our siLrif1 RNAseq data, where we did not detect increased expression from this class of repetitive elements (Figure S1A). Together, our results suggest an Lrif1-mediated regulation of *Dux* and Lrif1-independent regulation of IAPez repeats by Trim28.

Lastly, since the repressive histone modification, H3K9me3 is a known canonical marker of Trim28-mediated repression[22], we measured its levels at the *Dux* locus to test if reduced Trim28 binding at this locus upon Lrif1 knockdown leads to a concomitant decrease of this modification. ChIP-qPCR showed that both knockdown conditions resulted in decreased H3K9me3 levels at *Dux* (Figure 3F), however, this was attributable to the lower levels of H3 itself (Figure 3G) as the ratio of the modified H3 to all H3 remained unchanged (Figure 3H). This result is suggestive of increased chromatin accessibility at this locus and may explain the relatively subtle *Dux* expression changes upon knockdown of Lrif1 or Trim28 under these conditions.

Collectively, our findings identify a functional relationship between Lrif1 and Trim28 and support a conserved function of Lrif1 in the regulation of *Dux*/*DUX4* expression in mammals. It suggests that each of the FSHD2 genes currently identified, *SMCHD1, DNMT3B* and *LRIF1*, have distinct but perhaps partially overlapping roles in establishing and maintaining a repressive D4Z4 chromatin structure. How Lrif1 mediates *Dux* repression on a molecular level has to be further investigated. Alternative to direct binding to the *Dux* locus, an attractive possibility is that it acts through retinoic acid signalling, as LRIF1 was reported to be a nuclear receptor cofactor that interacts amongst others with retinoic acid receptors[14]. Indeed, retinoic acid is known to regulate the 2C-like cell transition [14, 24, 25].

## Data availability

The RNAseq datasets included in the study are available from GEO database under the accession number GSE249600. The mass spectrometry proteomics data have been deposited to the ProteomeXchange Consortium via the PRIDE [1] partner repository with the dataset identifier PXD042015.

## Acknowledgements

We thank Maja Vukic and Jihed Chouaref for their help with the RNAseq analysis and Dongxu Zheng for finalizing the RNAseq for resubmission. Our work is supported by the Prinses Beatrix Spierfonds (W.OR15-26) and the US National Institute of Arthritis and Musculoskeletal and Skin Diseases (R01AR066248). JB and SVDM are members of the European Reference Network for Rare Neuromuscular Diseases [ERN EURO-NMD].

## Declaration of interest

SMM is co-inventor on several FSHD patent applications and serves as a board member for Renogenyx. The other authors declare no competing interest.

## Web resources

OMIM, https://www.omim.org/

## Notes

https://www.ncbi.nlm.nih.gov/geo/

https://www.proteomexchange.org/

