## Supplemental figures, tables and methods for "Lrif1 modulates Trim28-mediated repression of the *Dux* locus in mouse embryonic stem cells"

**Figure S1**

A

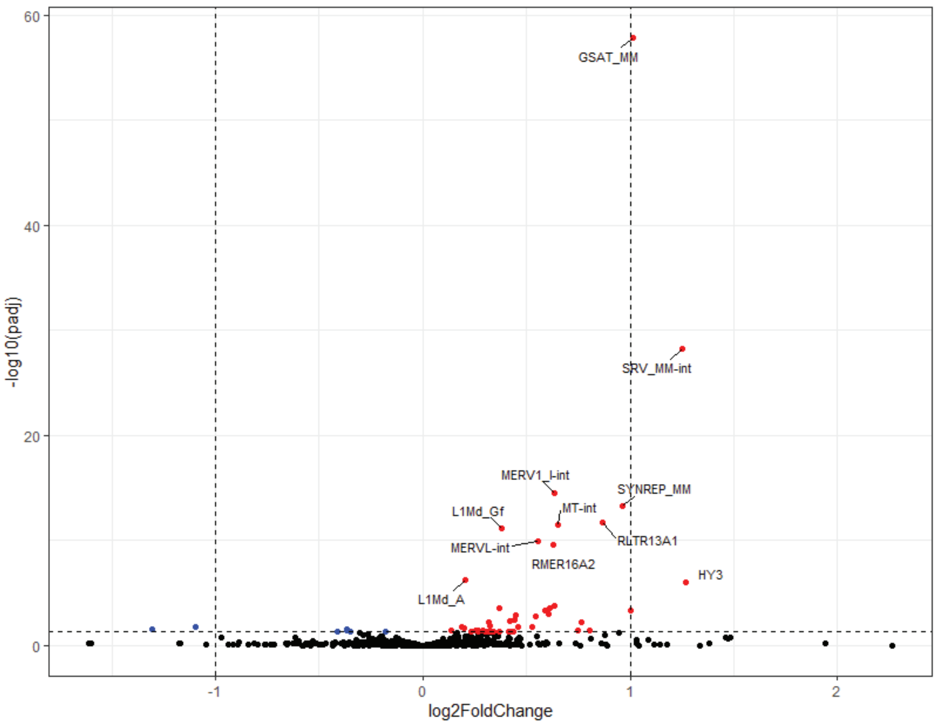


| DERs (padj < 0.05) | all | log2FC > 1 |
|---|---|---|
| ↑ repeats | 44 | 4 |
| ↓ repeats | 7 | 2 |

B

C

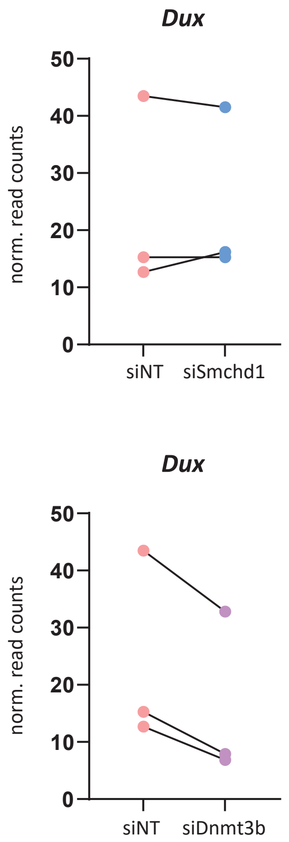


Figure S1. Lrif1 knock-down causes upregulation of 2C-specific repeats.
A) Volcano plot showing expression changes of repeats following Lrif1 knock-down. Upregulated repeats are highlighted in red and downregulated repeats are highlighted in blue. Dashed lines indicate a fold change of two (log2 fold of 1) on the x axis and significance of 0.05 (–log10 p.adj of 1.3) on the y axis. The top 10 differentially expressed repeats (DERs) are labelled. Table summary of DERs is provided below the plot. B) Normalized read counts of Dux transcripts after Smchd1 knock down compared to non-targeting siRNA condition. C) Normalized read counts of Dux transcripts after Dnmt3b knock down compared to non-targeting siRNA condition.

Figure S2

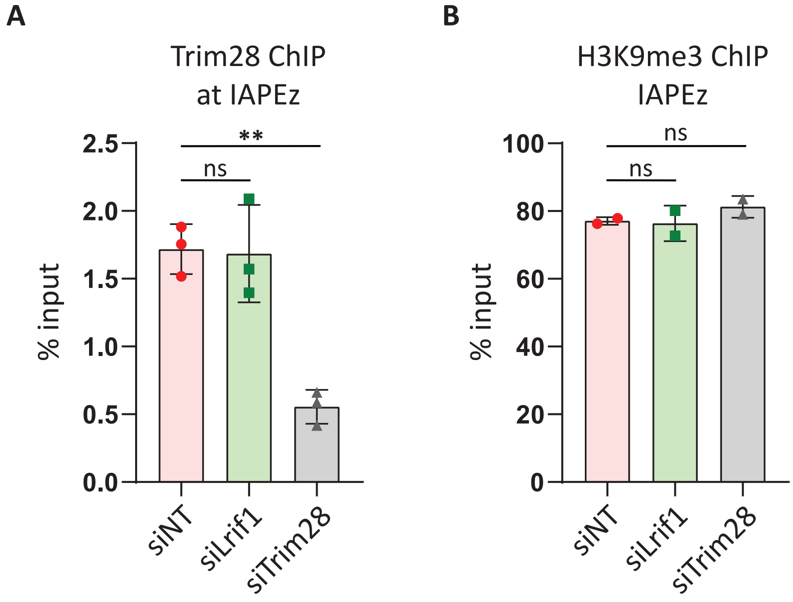


Figure S2. Lrif1 does not affect Trim28-mediated repression of IAPEz.

A) Trim28 ChIP-qPCR of the 5' IAPEz region in E14 mESCs after treatment with respective siRNAs. Bars and whiskers represent mean ± SEM of three independent experiments. Statistical significance was calculated by one-way ANOVA with 'Dunnett's post hoc test (ns: not significant; *: < 0.05, **: < 0.01). B) H3K9me3 ChIP-qPCR of the 5' IAPEz region in E14 mESCs after treatment with respective siRNAs. Bars and whiskers represent mean ± SEM of two independent experiments. Statistical significance was calculated by one-way ANOVA with 'Dunnett's post hoc test (ns: not significant).

**Supplemental Tables**

**Table S1.** Primers used for qRT-PCR.

| **Primer name** | **Primer sequence 5' ->3'** |
|---|---|
| mβ-actin_RT_F | GGCTGTATTCCCCTCCATCG |
| mβ-actin_RT_R | CCAGTTGGTAACAATGCCATGT |
| mLRIF1l+s qPCR F | AAGATGCAAACATTGTGGTG |
| mLRIF1l+s qPCR R | CCATCTTCATGGTTTCCGC |
| Smchd1 ex44_F | AAGCCCTTTGGAAATCCAGT |
| Smchd1 ex46_R | TGGGGCAGTGTGTGATTTTA |
| mDnmt3b_RT_Ex16-17_F | GGAAGAATTTGAGCCACCCA |
| mDnmt3b_RT_Ex18_R | GACTTCGGAGGCAATGTACTT |
| endo-mOct4-F | TAGGTGAGCCGTCTTTCCAC |
| endo-mOct4-R | GCTTAGCCAGGTTCGAGGAT |
| endo-mSox2-F | AGGGCTGGGAGAAAGAAGAG |
| endo-mSox2-R | CCGCGATTGTTGTGATTAGT |
| endo-mNanog-F | CTCAAGTCCTGAGGCTGACA |
| endo-mNanog-R | TGAAACCTGTCCTTGAGTGC |
| mTrim28-1241-F2 | CTGGTACGAACTCCACAGGT |
| mTrim28-1439-R2 | CCACTTACCTCTCCCTCACC |
| mDux_1 F | ACTTCTAGCCCCAGCGACTC |
| mDux_1 R | CCATGCTGCCAGGATTTCTA |
| Gm21761 F | GATCCCTGAGGGTAAGTCCTCC |
| Gm21761 R | TGCTTCCTATCCAGCTCTTGAGG |
| Usp17lb F | CTTCCCAGAAGATCCAGCC |
| Usp17lb R | CTGTGCTTTCCATTGGCAG |
| Gm2016 F | TACTCACCAGGTCAATGCAG |
| Gm2016 R | AGGAAGGTGTAGTCTCCCT |
| Tmem92 F | GTAAGCTTCAATGAGACTGCA |
| Tmem92 R | GCAGCATTCCTTGACACAG |
| mZscan4e-358-F | TTGAAGCCTCCTGTCATGGT |
| mZscan4e-515-R | TGTGTGGTGTCTACTGGCAT |

**Table S2.** Information about batch effects due to separate knockdown experiments and re-sequencing of the siDnmt3b_1 sample.

| Sample Name | Experiment ID | Sequencing run ID |
|---|---|---|
| siNT_1 | E1 | SR1 |
| siNT_2 | E2 | SR1 |
| siNT_3 | E3 | SR1 |
| siLrif1_1 | E1 | SR1 |
| siLrif1_2 | E2 | SR1 |
| siLrif1_3 | E3 | SR1 |
| siSmchd1_1 | E1 | SR1 |
| siSmchd1_2 | E2 | SR1 |
| siSmchd1_3 | E3 | SR1 |
| siDnmt3b_1 | E1 | SR2 |
| siDnmt3b_2 | E2 | SR1 |
| siDnmt3b_3 | E3 | SR1 |

**Table S3.** Cloning primers for creating mPVL and m1 mutants in Lrif1l/s ORF.

| Mutant Name | Insert | Forward primer (5'->3') | Reverse primer (5'->3') |
|---|---|---|---|
| Lrif1l mPVL | Insert 1 | CATGGTCCTGCTGGAGTTCGTG | GATGGTCAGGAATTCGAGTTTCACAGTCTCTCAAATCTTTAGTGAG |
| | Insert 2 | CTCACTAAAGATTTGAGAGACTGTGAAACTCGAATTCCTGACCATC | ACAGGGATTTCTTGTCTCCC |
| | Insert 1 + Insert 2 | CATGGTCCTGCTGGAGTTCGTG | ACAGGGATTTCTTGTCTCCC |
| Lrif1l m1 | Insert 1 | CATGGTCCTGCTGGAGTTCGTG | CTTTTTCTCTTAAAATTTGCTCAGCTTCTCTTATTTTTTCATCTCTGATG |
| | Insert 2 | CATCAGAGATGAAAAAATAAGAGAAGCTGAGCAAATTTTAAGAGAAAAAG | ACAGGGATTTCTTGTCTCCC |
| | Insert 1 + Insert 2 | CATGGTCCTGCTGGAGTTCGTG | ACAGGGATTTCTTGTCTCCC |
| Lrif1s mPVL | Insert 1 | CATGGTCCTGCTGGAGTTCGTG | GATGGTCAGGAATTCGAGTTTCACAGTCTCTCAAATCTTTAGTGAG |
| | Insert 2 | CTCACTAAAGATTTGAGAGACTGTGAAACTCGAATTCCTGACCATC | ACAGGGATTTCTTGTCTCCC |
| | Insert 1 + Insert 2 | CATGGTCCTGCTGGAGTTCGTG | ACAGGGATTTCTTGTCTCCC |
| Lrif1s m1 | Insert 1 | CATGGTCCTGCTGGAGTTCGTG | CTTTTTCTCTTAAAATTTGCTCAGCTTCTCTTATTTTTTCATCTCTGATG |
| | Insert 2 | CATCAGAGATGAAAAAATAAGAGAAGCTGAGCAAATTTTAAGAGAAAAAG | ACAGGGATTTCTTGTCTCCC |
| | Insert 1 + Insert 2 | CATGGTCCTGCTGGAGTTCGTG | ACAGGGATTTCTTGTCTCCC |

### Supplemental Methods

#### Cell culture

E14 mouse embryonic stem cells (mESCs) were grown on 0.1% gelatin (Sigma, #G-1890) coated plates on a UV-irradiated feeder layer of MEFs. E14 mESCs were maintained in medium composed of KnockOut™ DMEM (Gibco, #10829018) supplemented with 10% FBS (Biowest, #S1810), 1x MEM Non-Essential Amino Acids Solution (Gibco, #11140050), 2 mM L-Glutamine (Gibco, #25030149), 1 mM Sodium Pyruvate (Gibco, #11360070), 0.1 mM 2-Mercaptoethanol (Gibco, #31350010) and $10^5$ U/mL Leukemia Inhibitory Factor (EMD Millipore, #ESG1107). HEK293T cells were maintained in medium composed of Gibco DMEM, High Glucose, Pyruvate (Gibco, #119950) with addition of 10% FBS (Biowest, #S1810) and 1x Penicillin/streptomycin (Gibco, #15140122).

#### siRNA transfections

mESCs were reverse transfected with siGENOME siRNA SMARTpools (Horizon) at a final concentration of 40 nM using Lipofectamine™ RNAiMAX (Thermo Fisher Scientific, #13778030). siGENOME Non-Targeting Pool #2 was used as a negative control. Two days after the first transfection, cells were either harvested or 1/5$^{th}$ of the cells were reverse transfected again as per the first transfection. Cells were harvested for subsequent analysis two days post-transfection.

#### RNA isolation

Cells were harvested in Qiazol (Qiagen, #79306) and total RNA was isolated using RNeasy mini kit (Qiagen, #74101) accompanied by on-column DNase I treatment.

#### cDNA synthesis followed by RT-qPCR

cDNA synthesis was performed with poly-dT primers using the RevertAid H Minus First Strand cDNA synthesis kit (Thermo Fisher Scientific, #K1621). Gene expression was analyzed using iQ™ SYBR® Green Supermix (Biorad, #1708887) on a CFX384 Touch Real-Time PCR Detection System. All primers used for RT-qPCR are listed in Supplementary Table S3. mβ-actin was used as a housekeeping gene.

**RNA processing for total RNA-seq**

Total RNA was isolated as described above and RNA-seq was outsourced to Macrogen-Europe. Sequencing libraries were prepared with TruSeq Stranded Total RNA with Ribo-Zero Human/Mouse/Rat kit (supplier) according to the 'manufacturer's manual. Samples were sequenced as 100 bp paired-end on a HiSeq 4000 instrument.

**RNA-seq data analysis**

Quality assessment of the raw sequencing reads was done using FastQC v0.11.6. Adapters were removed by Trimmomatic v0.38 with parameters PE -threads 10 -phred33 ILLUMINACLIP:TruSeq3-PE.fa:2:30:10 LEADING:3 TRAILING:3 SLIDINGWINDOW:4:15 MINLEN:40. The remaining quality-filtered reads were aligned to the mouse reference genome mm10 with the corresponding annotation file from Ensemble using the STAR aligner v2.7.1 with parameters. Read count table was obtained with HTSeq-count v0.9.1 using the GENCODE MV23 annotation with the option ""–stranded reverse"". The differential expression analysis was done with DESeq2 v1.24.0 (R package). In the design we included correction for known experimental and sequencing batch effects (see Suppl. Table 4). Otherwise, default settings were used except for siSmchd1 analysis, where differential expression analysis was done on pre-filtered data, where 78% of the lowest expressed genes (based on row-wise mean of normalized counts) was filtered out due to otherwise faulty outcome when independent filtering was used. The final list of differential expressed genes contains genes for which the adjusted p-value (Benjamini-Hochberg correction) is < 0.05. RNA-seq plots were generated with ggplot2 package v3.3.3.

**Cloning**

The ORFs of Lrif1s and Lrif1l were subcloned first into pEF1a-FB-dCas9-puro (Addgene plasmid #100547) by replacing Cas9 insert via the NheI/XbaI sites. Lrif1s/l ORFs were amplified using an N-terminal specific primer for either Lrif1s (mLrif1s NheI F: 5'-TCCTTGCTAGCATGGCATCAATAGTAAAAAGGAAATTC-3') or Lrif1l (mLrif1l NheI F: 5'-

TCCTTGCTAGCATGTCTAATAGTCTCCAGAGCG-3') combined with common a reverse primer (mLrif1 XbaI R: 5'-CCTCATCTAGATTATTGTTTTTGGTACATCTTCTTACGC-3'). Resulting plasmids were further modified by replacing the N-terminal FlagAvi tag with EGFP via BstbI/NheI sites. EGFP was amplified from the pEGFP-C1 plasmid using the following primers: EGFP BstbI F: 5'-GATCTTTCGAAAGCCACCATGGTGAGCAAGGG-3' & EGFP NheI R:5'-ACATGCTAGCAAGGATCCTCGAAGCTTGAGCTCGAGATC-3'). A control plasmid expressing only EGFP-NLS was created by replacing the FlagAvi-Cas9 insert in pEF1a-FB-dCas9-puro with EGFP-NLS via the BstbI/XbaI sites. EGFP-NLS was amplified from the C1-EGFP-NLS plasmid (kind gift of Prof. Haico van Attikum, Leiden) with the following primers: EGFP BstbI F:5'-GATCTTTCGAAAGCCACCATGGTGAGCAAGGG-3' & EGFP-NLS XbaI R: 5'-CCTCATCTAGACTAAACCTTTCTCTTCTTCTTAGGACC-3'). The two C-terminal Lrif1 mutants (denoted in the text as mPVL and m1) were introduced to both the short and the long Lrif1 isoform and were created by overlapping PCR with mutagenic primers carrying the respective mutations (for primers combinations see Supplementary Table S3). Respective pEF1a-EGFP-Lrif1s/l-puro plasmids were used as a template for creating inserts 1 and 2, which were subsequently used in the final merging PCR. Final PCR products were cloned in the pEF1a-EGFP-Lrif1s/l-puro via BamHI/XbaI, thus exchanging the Lrif1s/l WT ORFs with their mutant counterpart. All vectors were sequence-verified by Sanger sequencing.

**Western blotting**

Samples were lysed with RIPA buffer (0.1% SDS, 1% Igepal CA-630, 150mM NaCl, 0.5% Sodium Deoxycholate, 20mM EDTA) supplemented with 1x Complete™, EDTA-free Protease Inhibitor Cocktail (PI) (Sigma-Aldrich, #11873580001). Samples were incubated on ice for 10 min followed by centrifugation at 15,000g for 15 min at 4°C. The protein concentration was determined with the Pierce™ BCA Protein Assay Kit (Thermo Fisher Scientific, #23225). For western blotting, samples were first mixed with 6X SB (0.375M Tris pH 6.8, 12% SDS, 60% glycerol, 0.6M DTT, 0.06% bromophenol blue) to 1x final concentration, boiled for 10 min at 95°C and resolved on Novex™ NuPAGE™ 4-12% Bis-Tris protein gels (Invitrogen, #NP0321BOX). Post-run the gel was transferred to an Immobilon-FL PVDF membrane (Merck, #IPFL00010). The membrane was blocked for 1 h in 4% skimmed milk in PBS followed by incubation overnight at 4°C with primary antibodies diluted in Immuno Booster solution I (Takara, #T7111A): RαGFP (1:1000, Abcam, #ab290), RαSMCHD1 (1:1000, Abcam #ab31865), RαLRIF1 (1:1000, Proteintech, #26115-1-AP), MαKAP1 (1:1000, Abcam, #ab22553), MαHP1γ clone 42s2 (1:1000, EMD Millipore, #05-690) and Mα-αTubulin (1:4000, Sigma-Aldrich #T6199). The next day, membranes were washed twice with PBS-T (0.01% Tween-20) and incubated with the following secondary antibodies diluted in Immuno Booster solution II (Takara, #T7111A): IRDye® 800CW goat anti-rabbit IgG (1:10,000, Li-cor #P/N 925-32211) and IRDye® 680CW donkey anti-mouse IgG (1:10,000, #P/N 925-68072) for 1h at room temperature. Membranes were washed twice with PBS-T prior scanning on an Odyssey® CLx Imaging System (Li-cor).

**Plasmids transfections**

For co-immunoprecipitation with GFP-trap beads, $10x10^6$ mESCs were reverse transfected with 5 µg of plasmid DNA with Lipofectamine 3000 (Thermo Fisher Scientific, #L3000008) on a Ø6 cm dish with 0.1% gelatine. Each transfection condition was done in three biological replicates. 30h post-transfection, cells were harvested for downstream co-immunoprecipitation with GFP-trap beads. For GFP co-immunoprecipitation in HEK293T cells, $2.5x10^6$ cells were seeded one day prior plasmid transfection on a Ø10 cm dish. The next day, cells were transfected with 6 µg of plasmid DNA with polyethylenimine (PEI) in a 1:3 volume ratio. Cells were harvested 30h post-transfection.

**GFP-Trap co-immunoprecipitation for mass spectrometry**

Transfected mESCs were washed 2x with ice-cold PBS and lysed on the dish with 600 µl of NP40 lysis buffer (50 mM Tris-HCl pH 8.0, 150 mM NaCl, 1% NP-40) supplemented with 1x PI, 20 mM NaF and 20 mM NEM. Whole cell lysates were incubated at 4°C, for 15 min while rotating, then spun down for

14,000g, 10 min at 4°C. 5% volume of supernatant was saved as input, mixed with 6xSB to a final 1X concentration and boiled for 10 min at 95°C. The remaining volume of the supernatant was added to 20 µl pre-washed GFP-Trap agarose beads (Chromotek, #gta-20) and incubated for 1.5 h at 4°C while rotating. Beads were subsequently washed 2x with NP40 lysis buffer followed by 3x wash with NP40 lysis buffer without NP40 and final three washes were done with freshly prepared 50 mM ammonium bicarbonate (ABC). After the last wash, 10% of the beads were used for protein elution in 2xSB by boiling for 15 min at 95°C while shaking to check for IP efficiency by Western blot. The rest of the beads were incubated over night with 2.5 µg of sequencing grade trypsin (Promega, #V5111) (dissolved in 50 mM ABC) at 37°C while shaking. The next day, digested peptides were filtered through a pre-washed 0.45 µm filter (EMD Millipore, #UFC40LH25) followed by acidification by addition of trifluoroacetic acid (TFA) to a 2% final concentration. Peptide solutions were loaded on custom-made Stage Tip as containing a disk made of a tC18 cartridge (Waters, #WAT036820) as described previously [16]. Stage Tips were washed twice with 0.1% formic acid, and peptides were eluted with 2x 25 µl of 32.5% acetonitrile in 0.1 % formic acid. Eluates were vacuum dried with a SpeedVac RC10.10 and kept at -80°C.

**Mass spectrometry data acquisition**

Mass spectrometry data was acquired essentially as in Gonzalez-Prieto et al. [17]. In brief, a liquid chromatography gradient was performed on an EASY-nLC 1000 system (Proxeon, Odense, Denmark) connected to a Q-Exactive Orbitrap (Thermo Fisher Scientific, Germany) through a nano-electrospray ion source. The Q-Exactive was coupled to a 20 cm analytical column with an inner-diameter of 75 µm, in-house packed with 1.9 µm C18-AQ beads (Reprospher-DE, Pur, Dr. Maish, Ammerbuch-Entringen, Germany). For each sample, two different acquisition methods were performed as technical repeats. The chromatography gradient length was 70 minutes from 2% to 30% acetonitrile in followed by 5 minutes gradient to 95% acetonitrile in 0.1% formic acid prior to column re-equilibration at a flow rate of 200 nL/minute. The mass spectrometer was operated in data-dependent acquisition (DDA) mode.

The first technical repeat was performed with a top-10 method. The maximum MS1 and MS2 injection times were 250 ms and 60 ms, respectively. For the second technical repeat, a Top5 method was used with MS1 and MS2 injection times were 250 ms and 256 ms, respectively. In both technical repeats, full-scan MS spectra were acquired in a range from 300 to 1600 m/z at a target value of $3 \times 10^6$ and a resolution of 70,000 and the Higher-Collisional Dissociation (HCD) tandem mass spectra (MS/MS) were recorded at a target value of $1 \times 10^5$ with a resolution of 17,500. Minimum AGC target was set to $1 \times 10^4$ and the normalized collision energy (NCE) was set to 25. Isolation window was 2.2 m/z wide. The precursor ion masses of scanned ions were dynamically excluded (DE) from MS/MS analysis for 20 sec. Ions with charge 1, and greater than 6 were excluded from triggering MS2 analysis.

**Mass spectrometry data analysis**

LC-MS/MS Raw files were analyzed using MaxQuant software (v1.6.14) according to Tyanova et al.[18] using default settings with the following modifications. Maximum number of mis-cleavages by trypsin/p was set to 3. Variable modifications included Oxidation (M), Acetyl (Protein N-term) and Phospho (STY) with a maximum number per peptide of 3. Carbamidomethyl(C) was deactivated as fixed modification. Label-free Quantification was enabled without the Fast LFQ algorithm. We performed the search against an in silico digested UniProt reference proteome for Homo sapiens (19th Sep 2019). Match-between-runs feature was enabled with 0.7 min match time window and 20 min alignment time window. Protein quantification included all the peptides. MaxQuant proteingroups.txt file output was further analyzed in Perseus computational platform (v1.6.14) as described by Tyanova et al.[18]. Potential contaminants and reverse peptides were removed, matrix was log2 transformed and proteins not identified in 3 out of 3 replicates for at least one condition were also removed. Missing values were randomly imputed from normal distribution width of 0.3 and a downshift of 1.8. Statistical conditions between groups were calculated by t-test with a permutation based FDR of 0.05 and an S0 of 0.1. Statistical tables were exported and data was further processed in Microsoft Excel 365 for comprehensive visualization.

**GFP-Trap co-immunoprecipitation with Lrif1 mutants in HEK293T cells**

Transfected HEK293T cells were washed 2x with ice-cold PBS and lysed on the dish with 1 ml of NP40 lysis buffer (50 mM Tris-HCl pH 8.0, 150 mM NaCl, 1% NP-40) supplemented with 1x PI, 20 mM NaF, 20 mM NEM, 2 mM $MgCl_2$ and 250U Benzonase (EMD Millipore, #E1014). Whole cell lysates were incubated at 4°C, for 1 h while rotating, then spun down for 14,000g, 10 min at 4°C. 1 mg of whole cell lysate was added to 20 µl pre-washed GFP-Trap agarose beads and incubated for 1.5 h at 4°C while rotating. Beads were washed 4x with NP40 lysis buffer and proteins were eluted by boiling the beads in 2xSB at 95°C for 15 min while shaking. Input samples represent 2% of material used for IP.

**Endogenous co-immunoprecipitation in mESCs**

mESCs were washed 2x with ice-cold PBS and lysed on the dish with EBC lysis buffers (50 mM Tris-HCl pH 8.0, 150 mM NaCl, 0.5% NP-40, 2 mM $MgCl_2$) supplemented with 1x PI, 20 mM NaF and 20 mM NEM. Whole cell lysates were incubated at 4°C, for 15 min while rotating, then spun down for 14,000g, 10 min at 4°C. 500 µg of whole cell lysate was added to 20 µl of antibody pre-linked Dynabeads and incubated over night at 4°C while rotating. Protein A Dynabeads (Thermo Fisher Scientific, #10002D) were used for conjugation of antibodies of rabbit origin and protein G Dynabeads (Thermo Fisher Scientific,#10003D) were used for Co-IP with antibodies of mouse origin. The following antibodies were used for endogenous Co-IPs: RαKap1 (Abcam, #ab10483), RαLRIF1 (Proteintech, #26115-1-AP), RαIgG (Cell Signalling, #2729S), MαKap1 (Abcam, #ab22553) and MαIgG (Merck, #12-371). The next day, beads were washed 4x with EBC lysis buffer and proteins were eluted by boiling the beads in 2xSB at 95°C for 15 min while shaking.

**Chromatin immunoprecipitation followed by qPCR**

Feeder MEFs were removed by pre-plating the trypsinized cell suspension 2x for 20 min on gelatinized culture plates. The supernatant was collected and washed once in 1x warm PBS followed by crosslinking with 1% formaldehyde in PBS for 10 min at room temperature while tumbling. The reaction was quenched by adding glycine to a final concentration of 125 mM. Crosslinked cells were washed twice with PBS, and the cell pellet was either stored at -80°C or proceeded to chromatin isolation. Cell pellets were resuspended in ice-cold ChIP buffer (1.5 ml lysis buffer/10 x $10^6$ cells) (150 mM NaCl, 50 mM Tris-HCl pH 7.5, 5 mM EDTA, 0.5 % Igepal CA-630, 1% Triton X-100) supplemented with cOmplete™ Protease Inhibitor Cocktail (Sigma-Aldrich, #11697498001). After 10 min incubation on ice, samples were spun down at 8,000 g for 2 min at 4°C. The pellets were resuspended for the second time in ChIP buffer, incubated for 5 min on ice and spun down again. Final nuclear pellets were resuspended in the ChIP buffer and sonicated at the highest power output for 15 cycles (1 cycle: 30 sec ON/30 sec OFF) using a Bioruptor instrument (Diagenode). For ChIP, chromatin was first pre-cleared with BSA-blocked protein A Sepharose beads (GE Healthcare, #17-5280-21) by rotating for 30-60 min at 4°C. For histone ChIP, 6 µg of chromatin was used; for Trim28 ChIP, 30 µg of chromatin was used in the final volume of 500 µl. 50 µl (10%) of each chromatin was kept as input sample for later normalization. ChIP was carried out by rotation at 4°C with following primary antibodies: RαTrim28 (Abcam, #ab10483), RαH3 (Abcam, ab1791), RαH3K9me3 (Active Motif, #39161) or rabbit polyclonal IgG (Abcam, #ab37415), which served as a negative control. Second day, 20 µl of protein A Sepharose beads pre-blocked with BSA were added to all samples and incubated for 2 h at 4°C while rotating. Afterwards, beads were washed as follows: once with low salt wash buffer (1 % Triton X-100, 0.1 % SDS, 2 mM EDTA, 20 mM Tris-HCl, 150 mM NaCl), high salt wash buffer (1 % Triton X-100, 0.1 % SDS, 2 mM EDTA, 20 mM Tris-HCl, 500 mM NaCl), LiCl wash buffer (250 mM LiCl, 1% Igepal CA-630, 1% sodium deoxycholate, 1 mM EDTA, 10 mM Tris-HCl) and twice with TE wash buffer (10 mM Tris-HCl, 1 mM EDTA). For DNA extraction, 10% (w/v) of Chelex 100 resin was added to the beads and boiled at 95°C for 15 min while vigorously shaking. The supernatant was used for qPCR analysis using the following primers for the Dux locus: Dux ChIP F2: 5’-CTAGCGACTTGCCCTCCTTCTG-'3' and Dux ChIP R2: 5’-ATTCAGAGGGGCTGGAGCAG-'3' at 60°C.
